# Hellbender Salamanders (*Cryptobranchus alleganiensis*) Exhibit an Ontogenetic Shift in Microhabitat Use in a Blue Ridge Physiographic Region Stream

**DOI:** 10.1101/139766

**Authors:** K. A. Hecht, M. J. Freake, M. A. Nickerson, P. Colclough

**Affiliations:** PO Box 116455, School of Natural Resources and Environment, University of Florida, Gainesville, Florida 32611;; Twitter: @HellbenderHecht; PO Box 117800 Florida Museum of Natural History, University of Florida, Gainesville, Florida 32611; Department of Natural Sciences and Mathematics, Lee University, Cleveland, Tennessee 37320; Zoo Knoxville, 3500 Knoxville Zoo Drive, Knoxville, Tennessee 37914

**Keywords:** Amphibian, Ecology, Lotic, Salamander, Shelter, Substrate

## Abstract

Organisms that experience large changes in body size during the life span often exhibit differences in resource use among life stages. Ontogenetic shifts in habitat use reduce intraspecific competition and predation and are common in lotic organisms. Although information on the immature life stages of the Hellbender (*Cryptobranchus alleganiensis*) is limited, this aquatic salamander exhibits ontogenetic shifts in habitat use in some streams, with adults sheltering under large rocks and larvae utilizing interstitial spaces of gravel beds. Due to the geomorphology of Little River, Tennessee, however, limited interstitial spaces within the gravel are filled with sand. Therefore, we quantified microhabitat parameters for three life stages of Hellbenders (larvae, sub-adult, adult) to determine if an ontogenetic shift in microhabitat occurred in Little River. We found no significant differences in stream substrate at capture sites among the stages, but there was a positive correlation between rock shelters underlain with very coarse gravel and overall Hellbender occupancy. Although we found no difference in water quality parameters and streambed particle size among the stage classes at the sites of capture, there was a significant difference in the average shelter size among all stages, with larvae utilizing the smallest shelters. Based on these results, future Hellbender research and conservation efforts should consider differences in life stage habitat use as well as specific stream particle classes.

---

Body size is a key factor in many facets of ecology. At larger scales, the size of species helps determine the trophic structure and spatial distribution of ecological communities (Hutchinson and MacArthur, 1959; Schoener, 1974; Werner and Gilliam, 1984; Brown and Nicoletto, 1991; Woodward et al., 2005; Rojas and Ojeda, 2010), while at the individual scale body size influences energetics (Gillooly et al., 2001), prey (Wilson, 1975; Mittelbach, 1981; Cohen et al., 1993), habitat use (Hall and Werner, 1977; Foster et al., 1988; Flinders and Magoulick, 2007; Barriga and Battini, 2009; Foster et al., 2009), and predation risk (Werner and Hall, 1988; Giller and Malmqvist, 1998; Urban, 2008). Because size has such a strong influence on the ecology of organisms, species that experience large changes in body size during their lifespan can experience substantial differences in ecology across life stages. Werner and Gilliam (1984) defined these changes (i.e., ontogenetic shifts) as the “patterns in an organism’s resource use that develop as it increases in size from birth or hatching to its maximum.” While these changes are often a result of morphological constraints, change in resource use across the life span of a species can be an advantageous life history strategy. These shifts may reduce intraspecific competition and predation among stage classes (Werner and Gilliam, 1984). In cannibalistic species shifts in habitat use among size or stage classes can reduce mortality of young individuals by intraspecific predation (Foster et al., 1988; Keren-Rotem et al., 2006).

Body size changes in species are especially relevant in lotic systems. Reynolds number, which is the ratio of inertial and viscous forces within a fluid, increases with body size (Giller and Malmqvist, 1998). Organisms with different Reynolds numbers experience varying impacts from stream flow with inertial forces becoming more important at higher Reynolds numbers and may also differ in gas exchange abilities (Giller and Malmqvist, 1998). Body size influences microhabitat use in streams, with larger individuals more likely to reside in the water column and smaller animals governed by viscous forces typically inhabiting the stream substrate. Because of these differences, ontogenetic shifts in resource use are documented in aquatic organisms and occur in a wide range of lotic taxa across different trophic levels including invertebrates (Holomuzki and Short, 1990; Giller and Sangpradub, 1993; Flinders and Magoulick, 2007), fish (Merigoux and Ponton, 1998; Simonovic et al., 1999; Rosenberger and Angermeier, 2003; King, 2005; Barriga and Battini, 2009) and salamanders (Petranka, 1984; Colley et al., 1989; Nickerson et al., 2003). These shifts in resource use among life stages may help mitigate challenging conditions in lotic environments such as flow, environmental variability, and limited dispersal potential by providing increased protection and food availability and decreased intraspecific competition (Werner and Hall, 1988; Colley et al., 1989; Giller and Malmqvist, 1998; Nickerson et al., 2003; Barriga and Battini, 2009)

Ontogenetic shifts in resource use have been noted in the Hellbender (*Cryptobranchus alleganiensis*), a cannibalistic lotic salamander species that can increase in size over its lifetime by a factor of 20. Hatchlings measure 25 – 30 mm total length (TL), while the largest adult found measured 745 mm TL (Fitch, 1947). Larval Hellbender diet largely consists of aquatic insects (Smith, 1907; Pitt and Nickerson, 2006; Hecht et al., 2017) whereas adults mostly eat crayfish (Netting, 1929; Green, 1933; Green, 1935; Nickerson and Mays, 1973; Peterson et al., 1989). While there are very little data available on larval Hellbender ecology due to a lack of captures during surveys, researchers have noted that larval Hellbenders in some localities can utilize different microhabitat than adults, which generally shelter under large rocks (Bishop, 1941; Hillis and Bellis, 1971; Nickerson and Mays, 1973, Freake and DePerno, 2017). In the North Fork of the White River, Missouri, larvae have been associated with gravel beds (Nickerson et al., 2003), whereas bank searches in the Allegheny River, New York, were more effective for smaller Hellbender size classes than in previous conventional rock lifting surveys (Foster et al., 2009).

In Little River, Tennessee geology of the streambed led to sand and other small particles filling in the interstitial spaces within the gravel where larvae have been found in other streams (Nickerson et al., 2003; Pitt et al., 2016); thus, larvae have been found under rocks on the streambed surface like adults (Nickerson et al., 2003). Despite this difference, almost a third of sampled Hellbenders from Little River were larval sized (<125 mm) (Hecht-Kardasz et al., 2012). Due to the cannibalistic nature of Hellbenders (Humphries et al., 2005; Groves and Williams, 2014) as well as the great change in size from hatching to maturation, we expected that Hellbenders would still exhibit ontogenetic shifts in microhabitat at this location. To test this hypothesis, we examined the following microhabitat factors at sites where we captured Hellbenders in Little River: water depth, shelter size, stream substrate, pH, conductivity, and water temperature. These factors are known to affect detectability, food sources, oxygen concentration, and health of aquatic organisms (Giller and Malmqvist, 1998).

## MATERIALS AND METHODS

### Site description

Based on the results of a previous study (Nickerson et al., 2003), Hellbender surveys were conducted within an ~3 km protected and forested section of Little River known to contain the three stage classes (larvae, sub-adult, and adult). Little River, located in the eastern Tennessee portion of the Great Smoky Mountains National Park, originates on the north slope of Clingmans Dome, and flows 29 km within the park. It continues through the towns of Townsend, Maryville, Alcoa, and Rockford before eventually draining into the Tennessee River. The Little River watershed drains an area of approximately 980 km^2^.

Little River lies entirely within the southern portion of the Blue Ridge physiographic province. The bedrock of Little River is comprised primarily of late Precambrian Elkmont and Thunderhead metamorphosed sandstone (Mast and Turk, 1999). Over time flowing water has eroded away some exposed bedrock leaving large densities of rounded boulders, cobble, and gravel in the streambed. A Wolman pebble count (Wolman, 1954) in the study area found a D50 value, which represents the median substrate size, in the very coarse gravel category (32--64 mm) (Hecht-Kardasz, 2011). Interstitial habitat is limited within the Little River streambed as sand often fills in many portions of the gravel beds. The elevation of the study area ranged from 327— 407 m. Vegetation within the stream was uncommon, and the riparian vegetation was classified as pine and river cove hardwood forest (Madden et al., 2004). The area has a temperate climate, with an average annual rainfall of 142 cm and temperature averages of 3.17 °C in winter and 21.7 °C in summer (National Oceanic and Atmospheric Administration, 2016).

### Field methods

Diurnal skin diving combined with rock lifting was used to survey for Hellbenders during the following sampling periods: June – July 2005, June – July 2006, June – Aug 2008, Aug – Oct 2009, July – Sept 2010. Some surveyors occasionally used log peaveys to lift larger rocks. Hellbenders were captured by hand. We measured total length (TL) and snout-vent length (SVL) of most sub-adult and adult Hellbenders with the aid of modified PVC pipe. Hellbenders were individually marked before release using PIT tags. Larvae and sub-adults too small for PIT tags were marked using visible implant elastomer (see Hecht-Kardasz et al., 2012). We only included the initial habitat data from recaptured animals for analyses.

Microhabitat parameters were measured directly at the point of capture. Because Hellbenders are largely nocturnal (Nickerson and Mays, 1973) and generally have small home ranges and exhibit site fidelity (Hillis and Bellis, 1971; Wiggs, 1977; Nickerson and Mays, 1973; Blais, 1996; Ball, 2001), we assumed that the microhabitat at point of capture accurately represented microhabitat of Hellbenders during the survey period. Water temperature, pH, and conductivity were measured using the Combo pH/EC/TDS/Temperature Tester with Low Range EC and Watercheck pH reader (HANNA Instruments®, Woonsocket, RI, USA). Water depth and shelter size, defined as the longest length of the shelter rock, were also recorded.

To test for differences in stream substrate associated with shelter rocks, we measured a handful of streambed particles under confirmed shelter rocks using the Federal Interagency Sedimentation Project (FISP) US SAH-97 sediment size analyzer, also known as a gravelometer. Samples ranged from 1 – 8 particles, with a mean of 4.23 (± 1.55) particles. To compare the stream substrate beneath shelters with the streambed particles in the general sampling area, we also measured a handful of substrate at fifty random localities within the study area chosen using a random number table. Samples were taken directly next to the right foot with eyes averted. We sampled below larger rocks when they were encountered.

### Analyses

Individual Hellbenders were classified into stage classes using TL. We used TL in our analyses so we could directly compare our results to past Hellbender habitat studies (Hillis and Bellis, 1971; Humphries and Pauley, 2005). Individuals <125 mm in TL, both gilled and non-gilled, were classified as larvae. Larvae were also classified into first (<90 mm TL) and second year (>100 mm TL) age classes for shelter size analysis based on previous studies and the results of surveys in Little River (Smith, 1907; Bishop, 1941; Hecht-Kardasz et al., 2012). Three individuals between 90 – 100 mm TL could not be classified to an age class and were therefore not used in analysis comparing larval age classes. All individuals measuring 125 – 275 mm TL were considered sub-adults, while any individuals over 275 mm were classified as adults. Further justification for stage class classifications can be found in Hecht-Kardasz et al., 2012.

We analyzed data using base packages in R version 3.2.2 (R Core Team, 2015) unless otherwise specified. We calculated mean (+ SD) for all continuous normally-distributed habitat variables and median for non-normal continuous variables. Pearson’s correlation coefficients for all variables was below 0.5. To examine the relationships between habitat variables and Hellbender TL, we performed simple linear regressions. Habitat parameters were also compared among life stages. As water depth, larval shelter size, and conductivity data were not normally distributed, these parameters were tested using Kruskal-Wallis rank sum tests with pairwise comparisons performed using the pairw.kw function in the asbio package (Aho, 2014). The remaining normally distributed parameters were evaluated using ANOVA and t-tests. In order to control family wise error rate at 0.05, Bonferroni’s correction was used for all individual pairwise test of means.

All streambed particle sizes were classified into categories according to the American Geophysical Union proposed grade scale (Lane, 1947). Due to the low presence of some categories, all particles <4 mm were combined into one category before the data were used for statistical analysis. The presence/absence of streambed particle size at the site of capture was compared among stage classes using an ordinal logistic regression with the lrm function in package rms (Harrell, 2015). We also performed a binary logistic regression model using the lrm function to compare the presence/absence of particle categories between occupied sites and random locations. Due to weak correlations between smaller streambed particle size categories, additional models were tested combining all particles <32 mm into one category.

## RESULTS

Average pH at capture sites was 7.24 + 0.28 (range 6.74 – 8.10; n = 97). Mean conductivity was 12.98 + 2.41 μS/cm (range: 6.00 – 22.00 μS/cm; n = 79). Water depth (range: 210 – 1800 mm; n = 104) and water temperature (range: 14.60 – 22.80 °C; n = 103) averaged 527.86 + 248.00 mm and 22.84 + 2.03 °C respectively. Although regression analysis suggested a linear relationship between Hellbender TL and water temperature (n = 102), water temperature was not a strong predictor of Hellbender TL (R^2^ = 0.042; p = 0.039). A similar relationship was found between conductivity and Hellbender TL (R^2^ = 0.080; p = 0.012; n = 78). Linear regression analysis revealed no relationship between Hellbender TL and water depth (n = 104) (R^2^ = 0.024; p = 0.12) or Hellbender TL and pH (n = 96) (R^2^ = −0.011; p = 0.94). No significant difference in average water depth (H(2) = 4.32; p = 0.12), pH (F(2,97) = 0.61; p = 0.55) or temperature (F(2, 99) = 1.751; p = 0.179) was found among stage classes. Average conductivity was significantly different among stage classes (H(2) = 8.03; p = 0.018). Posthoc pairwise comparisons found a significant difference between larval mean conductivity (14.93 + 4.34 μS/cm; n = 14) and mean adult conductivity (12.53 + 1.59 μS/cm; n = 43; p = 0.018). There was no significant difference between larval and mean sub-adult conductivity (12.59 + 1.30 μS/cm; n = 22; p = 0.051) or between adult and sub-adult conductivity (p = 0.99) (Fig. 1).

**Figure 1.**
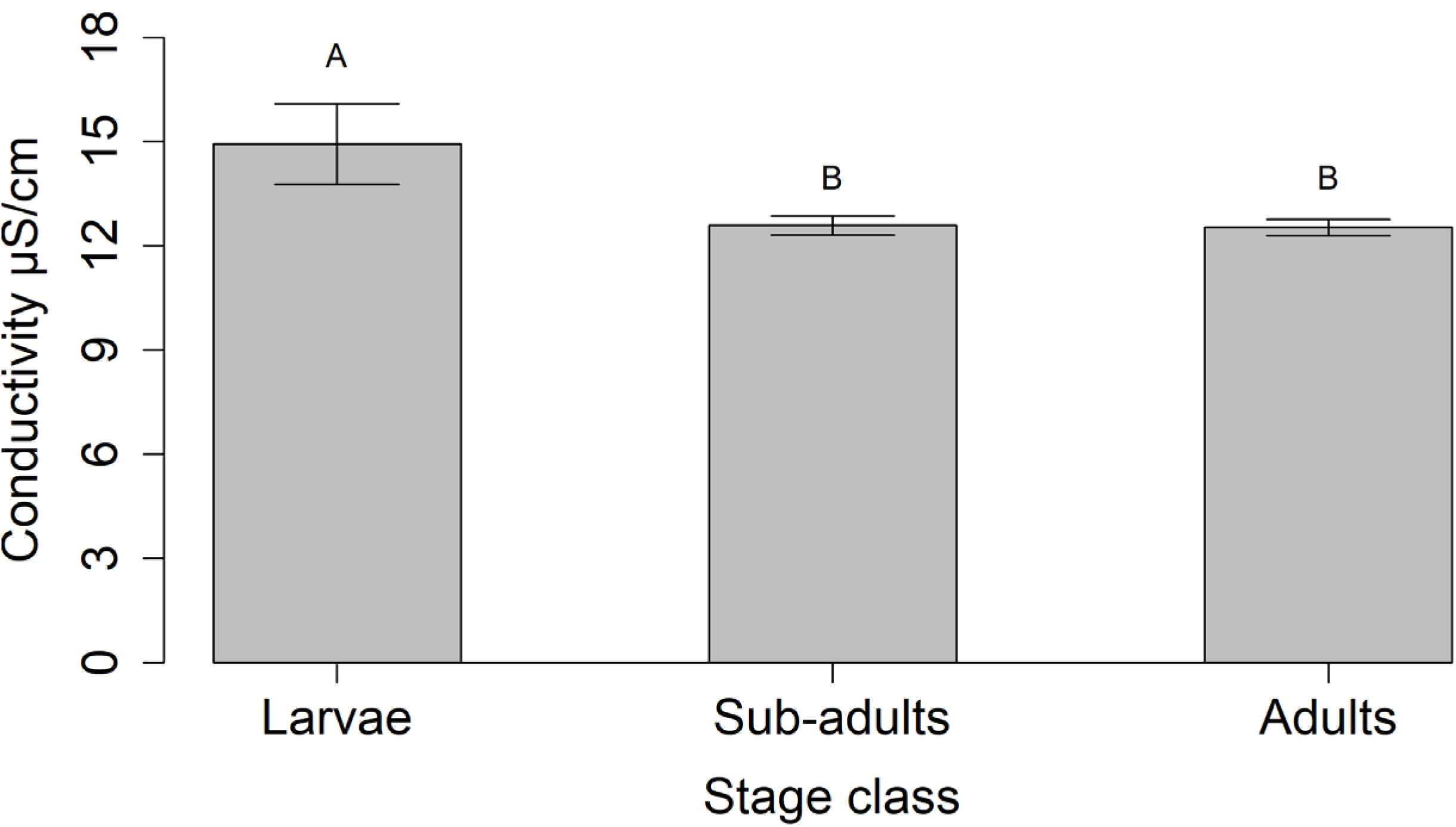
Bar graph showing mean ± standard error of the mean (SEM) for conductivity (μS/cm) used by three stage classes of *Cryptobranchus alleganiensis*, larvae (n = 13), sub-adults (n = 22), and adults (n = 43), in Little River, Tennessee. Bars with different letters above are significantly different (p < 0.05).

Shelter size ranged from 120 – 1470 mm with a mean of 673.81 + 285.75 mm (n = 217). Based on the results of linear regression, we found a weak correlation between Hellbender TL and shelter size (n = 217) (R^2^ = 0.266; p < 0.001) (Fig. 2). Although overall shelter size among the stage classes overlapped, average shelter size differed significantly among stage classes (F(2, 214) = 32.82; p < 0.001; Fig. 3). Mean shelter size of larvae (464.36 + 244.65 mm; n = 61) was significantly different from both adults (794.44 + 254.27 mm; n = 100; t = 8.11, df = 159, p-value = <0.001) and sub-adults (686.55 + 252.46 mm, n = 56; t = −4.83, df = 115, p = <0.001). Sub-adults (n = 56) and adults (n = 100) also differed significantly in mean shelter size (t = 2.55, df = 154, p = 0.012). There was no statistical difference between mean shelter size between first (n = 49) and second year larvae (n = 9) in Little River (H(1) = 0.16, p = 0.69). However, first year larvae utilized some larger shelter sizes, including one of 1085 mm while the largest shelter size of second year larvae was 610 mm. One individual of 90 mm TL found beneath a 1286 mm boulder could not conclusively be categorized as a first or second year larva.

**Figure 2.**
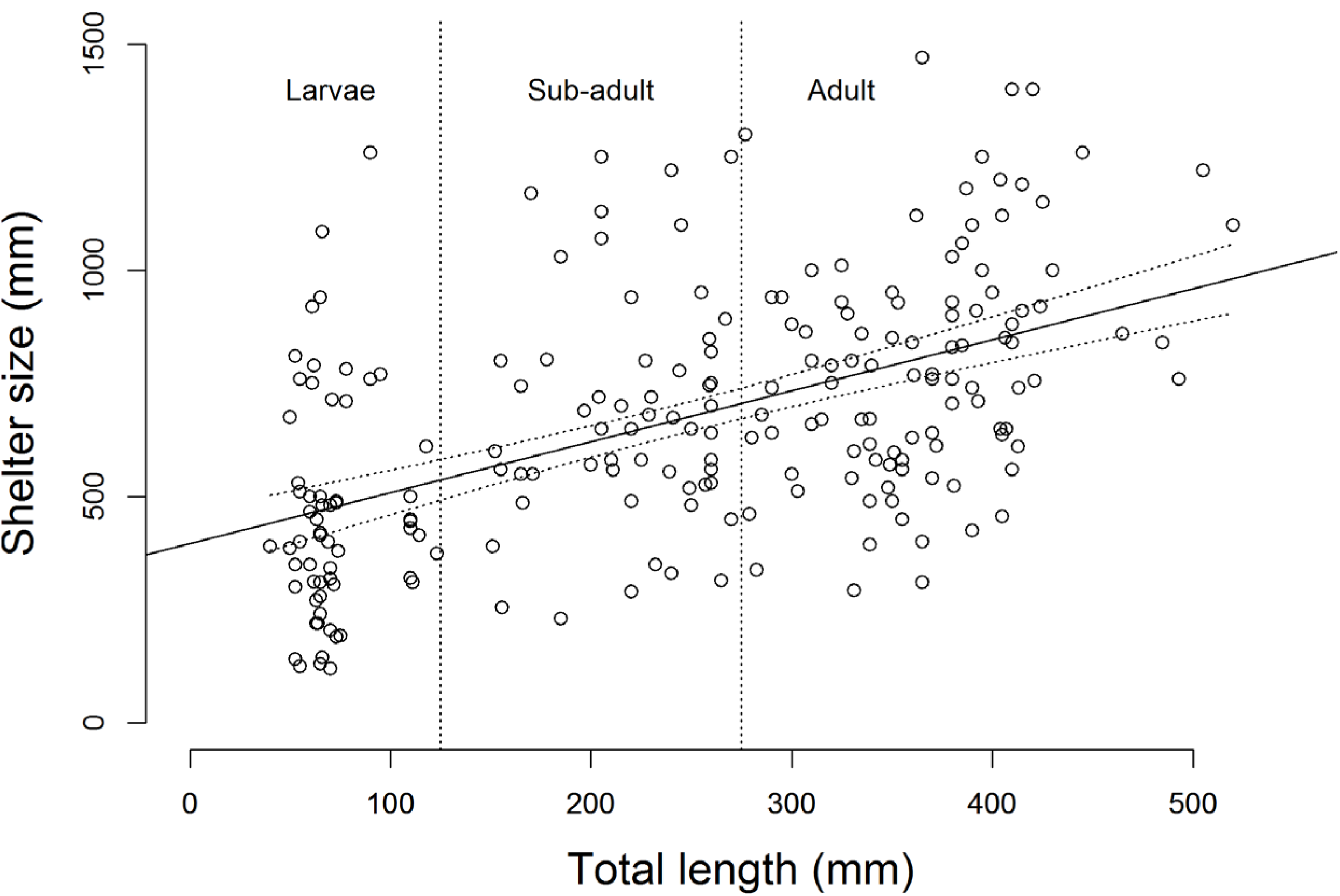
Scatter plot with linear regression line of shelter size (mm) vs. *Cryptobranchus alleganiensis* total length (mm) in Little River, Tennessee (n = 217) (R^2^ = 0.266; p<0.001).

**Figure 3.**
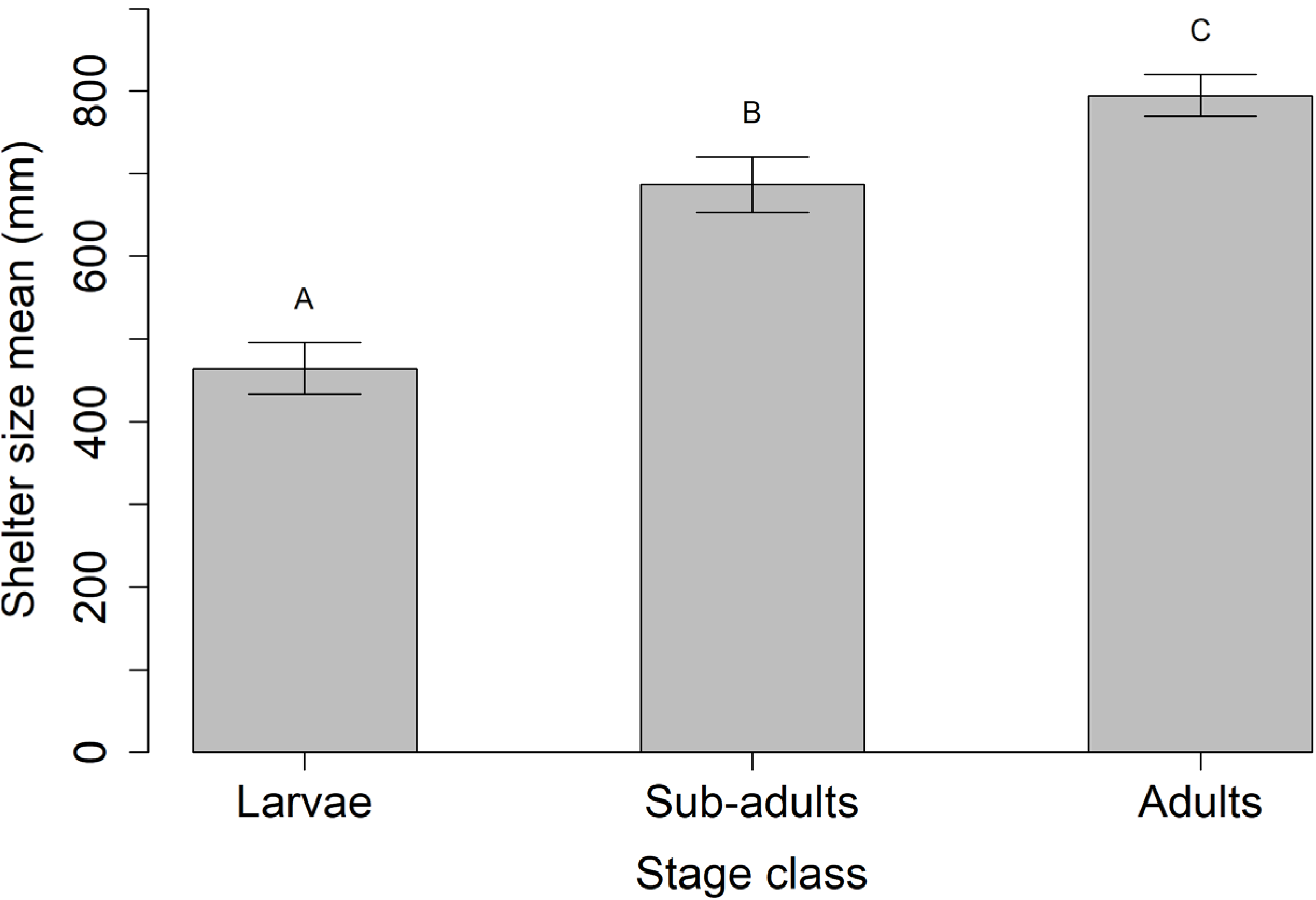
Bar graph showing mean ± standard error of the mean (SEM) for shelter size (mm) used by three stage classes of *Cryptobranchus alleganiensis*, larvae (n = 61), sub-adults (n = 56), and adults (n = 100), in Little River, Tennessee. Bars with different letters above are significantly different (p < 0.05).

Streambed particle classes under shelter rocks of larvae (n = 25), sub-adults (n = 26), and adults (n = 38) did not differ significantly (Table 1). There was no significant difference when particles <32 mm were combined. When comparing random samples to locations of capture, however, Hellbenders appeared to utilize shelters underlain at least partially by very coarse gravel more than would be expected by chance (Table 2). Our model also found a negative association between Hellbender use and rock shelters overlaying fine gravel. Very coarse gravel was the only significant term in the model combining particles <32mm (p < 0.001).

**Table 1.** Variable estimates and odds ratios from an ordinal logistic regression model based on streambed particle size classes at sites used by larval (n = 25), sub-adult (n=26), and adult (n=38) Hellbenders (*Cryptobranchus alleganiensis*) captured in Little River, Tennessee.

| Variable | Estimate | Standard<br>error | Wald statistic (Z) | p-value | Odds<br>ratio |
| --- | --- | --- | --- | --- | --- |
| <4 mm | 1.09 | 1.36 | 0.80 | 0.43 | 2.96 |
| Fine gravel | 0.66 | 1.13 | 0.58 | 0.56 | 1.93 |
| Medium gravel | -0.39 | 0.54 | -0.73 | 0.47 | 0.68 |
| Coarse gravel | -0.23 | 0.48 | -0.48 | 0.62 | 0.79 |
| Very coarse gravel | 2.13 | 1.20 | 1.78 | 0.07 | 8.45 |
| Small cobble | -0.54 | 0.46 | -1.19 | 0.23 | 0.58 |
| Large cobble | -0.52 | 0.49 | -1.06 | 0.29 | 0.59 |

**Table 2.**
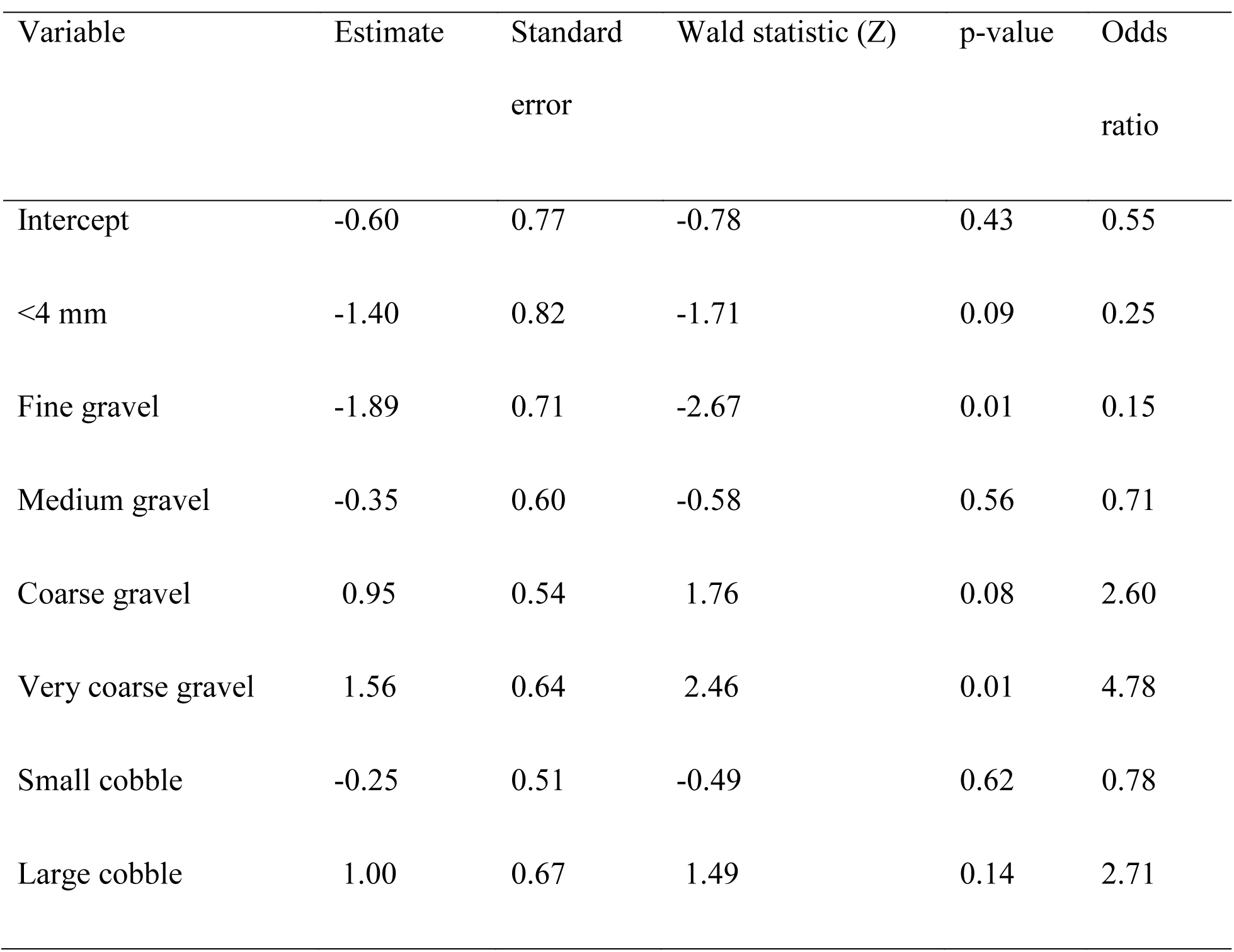
Variable estimates and odds ratios from a binomial logistic regression model based on streambed particle size classes at sites used by Hellbenders (*Cryptobranchus alleganiensis*) (n = 89) and random locations (n = 50) within Little River, Tennessee.

| Variable | Estimate | Standard<br>error | Wald statistic (Z) | p-value | Odds<br>ratio |
| --- | --- | --- | --- | --- | --- |
| Intercept | -0.60 | 0.77 | -0.78 | 0.43 | 0.55 |
| <4 mm | -1.40 | 0.82 | -1.71 | 0.09 | 0.25 |
| Fine gravel | -1.89 | 0.71 | -2.67 | 0.01 | 0.15 |
| Medium gravel | -0.35 | 0.60 | -0.58 | 0.56 | 0.71 |
| Coarse gravel | 0.95 | 0.54 | 1.76 | 0.08 | 2.60 |
| Very coarse gravel | 1.56 | 0.64 | 2.46 | 0.01 | 4.78 |
| Small cobble | -0.25 | 0.51 | -0.49 | 0.62 | 0.78 |
| Large cobble | 1.00 | 0.67 | 1.49 | 0.14 | 2.71 |

## DISCUSSION

While all Hellbender stage classes utilized boulder habitat, the significant difference in average shelter size among stage classes suggests that an ontogenetic shift in Hellbender habitat use occurs in Little River during the summer months. However, the wide range of shelter sizes used by larvae includes a direct overlap in shelter size with sub-adults and adults, which may be partially due to some young individuals dispersing from their site of hatching later than others. Young Hellbenders may remain in nesting sites for prolonged periods, as larval Hellbenders have been observed sharing rock shelters with adult males in June and August (Groves et al., 2015). Second year larvae could be more selective in their choice of shelter due to experience with predators, however the sample size of second year larvae was relatively small so further research is warranted. The weak relationship of shelter size and Hellbender TL found during this study is notable because previous studies examining habitat use by Hellbenders have generally found no association between shelter size and Hellbender size (Hillis and Bellis, 1971; Humphries and Pauley, 2005). However, these studies have focused primarily on adult-sized Hellbenders. A study in a 350 m section of the dam-impacted Hiawassee River (TN) found a similar pattern of shelter size use in a broader representation of Hellbender size classes (Freake and DePerno, 2017).

Flooding has been cited as a potential threat to Hellbender populations with several published reports of displaced, injured, and dead Hellbenders following high water events in other localities (Humphries, 2005; Miller and Miller, 2005; Bodinof et al., 2012a). Previous work in Little River suggested that flooding may be influential in the size structure of the Hellbender population with anecdotal evidence showing absent size classes correlating with major flooding events (Nickerson et al., 2007; Hecht-Kardasz et al., 2012). The shelters used by immature Hellbenders in Little River could provide a mechanistic explanation for this hypothesis. Many lotic organisms survive spates by seeking refugia (Giller and Malmqvist, 1998), including the interstitial spaces in the benthic layers, where larval *C. alleganiensis* have been located in other localities (Smith, 1907; Nickerson and Mays, 1973; Nickerson et al., 2003). As this habitat is not available to larval Hellbenders in Little River, larvae are utilizing the space under rocks at the surface of the streambed which may be less secure during flooding periods. While larvae utilized a wide variety of shelters in Little River, their habitat included much smaller shelter sizes than other stage classes including small and large cobble, and the average shelter size used by larvae was significantly smaller than sub-adults and adults. Smaller shelters may be easily moved by increased water current, increasing the risk of the Hellbender larvae underneath being crushed, swept downstream, or exposed to predators. Researchers recently found a crushed larvae in Little River following a high water event (Da Silva Neto et al., 2016). Related mortality or displacement of immature Hellbenders during extreme flooding related to less secure habitats may partially be responsible for the size structure patterns previously found in Little River’s captured Hellbender population (Hecht-Kardasz 2012). As increases in flood intensity and frequency are predicted with climate change (Easterling et al., 2000), this could be of conservation concern for Hellbenders, particularly in rivers with similar geomorphology although additional study is required.

Due to the lack of gravel bed habitat in Little River, the interstitial spaces among the gravel, cobble, and boulders beneath the larger shelter rocks may be particularly important to Hellbender larvae for additional protection and access to smaller food items. However, larvae were found directly under shelter rocks rather than underlying cobble or gravel (Hecht, pers. obs), and no difference in stream particle sizes below shelter rocks was noted among the stage classes. This suggests that other factors might be influencing habitat selection by Hellbenders in relation to substrate beneath shelter sites. For example, Bodinof et al (2012b) found that spacing of substrate was an important factor in Hellbender habitat selection for released captive raised Hellbenders, with individuals being more likely to select habitat resources where coarse substrate was touching.

Comparing streambed particle sizes at sites utilized by Hellbenders of all stage classes to randomly sampled localities revealed a negative association of occupancy with fine gravel, and a positive association of occupancy with very coarse gravel. It is unclear if these associations are due to habitat preferences and/or prey availability, or are simply related to space availability beneath shelter rocks. Smaller streambed particles could fill in the spaces underneath rocks, embedding them and leaving no area available for Hellbenders to occupy. Stream embeddedness has been negatively associated with the presence of other species of salamanders (Tumlinson and Cline, 2003). Conversely, boulders or large cobble may leave too much space available beneath shelter rocks, leaving Hellbenders with reduced protection from stream flow, predators, and con-specifics. The association of shelters used by Hellbenders and medium-sized particles, like very coarse gravel, may represent a balance of space availability and protection as well as food availability. Other studies have examined the role of streambed particle sizes on the occupancy of Hellbenders (Keitzer et al., 2013; Maxwell, 2009; Burgmeier et al., 2011; Bodinof et al, 2012b) but have been unable to compare streambed particle association among stage classes. Most of these studies have focused on broader particle categories rather than the more fine scale categories used in this study, but have found a general association between gravel and/or cobble substrates and Hellbender occupancy. These types of streambed particles are known to harbor a number of salamander species including Hellbender larvae (Smith, 1907; Nickerson and Mays, 1973; Tumlinson et al., 1990) and also serve as important macro-invertebrate habitat (Giller and Malmqvist, 1998; Hwa-Seong and Ward, 2007), which represent the most utilized food source for Hellbenders of all sizes.

Conductivity at larval sites was significantly different from adult sites. As conductivity measurements were low, and because there was little difference between the mean of the larval and other stage groups, it seems unlikely that this difference is biologically meaningful. However, conductivity impacts Hellbender distribution in other localities (Keitzer et al., 2013; Pitt et al., 2017; Bodinof Jachowski and Hopkins, 2018). No other correlations between Hellbender TL or stage class and measured water quality parameters were noted. The majority of individuals in all three stage classes were found in runs, so mixing may have created largely homogenized water quality conditions. Parameters including pH and conductivity showed little temporal or spatial variation during the survey period, but as Little River is fed by surface water, water depth and water temperate varied due to fluctuations in precipitation. Because microhabitat parameters were assumed to be relatively constant through time, this study cannot conclusively rule out the effects of water depth and water temperature on ontogenetic habitat use during the survey period.

Our examination of Hellbender microhabitat associations assumed that individuals were associated with the microhabitat at diurnal capture sites for significant time periods, and Hellbenders had similar detection rates across stage classes. While most studies support an association of adult Hellbenders to seasonal or longer habitats (Smith, 1907; Green, 1933; Hillis and Bellis, 1971; Wiggs, 1977; Nickerson and Mays, 1973; Nickerson, 1980; Blais, 1996; Ball, 2001), information regarding detectability, movement, activity, and site fidelity of immature Hellbenders is extremely limited. We are not aware of any studies available examining detection rates of immature Hellbenders. Since we did not find other available habitat types like gravel beds and leaf litter in the study sites and regularly located larval and sub-adult Hellbenders, we assumed that detectability rates were roughly the same among stage. Published information on larval movement is limited to a single observation of an individual moving along the stream margin an hour before sunset (Floyd et al., 2013). It is unclear whether *C. alleganiensis* larvae are nocturnal or diurnal in the wild, although Smith (1907) noted that hatchlings avoided light. Although it is also unknown whether wild Hellbender larvae leave shelter to forage, other salamander larvae have reduced activity levels in the presence of predators, including cannibalistic conspecifics (Colley et al., 1989). In addition macro-invertebrates found in larval Hellbender diets are plentiful beneath rocks in Little River (Hecht-Kardasz, 2011), thus low larval Hellbender activity might be expected. Larvae overwinter at male-guarded nest sites, and are believed to generally disperse sometime in spring or early summer (Bishop, 1941), prior to the seasonal timeframe of this study. As we already discussed above, some larvae may leave nest shelters later in the summer, but those captured during this study were almost entirely solitary, making it likely that dispersion had already occurred. While it is not unreasonable to assume that young Hellbenders, like adults, are associated with specific locations for extended periods and that detection rates were similar among the stage classes, these assumptions cannot be confirmed, and therefore the results of the analyses presented here should be interpreted with caution.

Evidence is increasing that Hellbenders may exhibit ontogenetic shifts in habitat use, but the number of localities where larval individuals are found regularly is relatively small, making it difficult to determine how common this pattern may be across the range. Other streams may have low larval detection rates making it more difficult to locate and quantify larval habitat. Future tracking of larvae may help elucidate whether larvae are rare or are avoiding detection due to differences in microhabitat use. In addition, only a limited number of microhabitat parameters have been examined. Therefore, studies looking at additional parameters such as DO, stream flow, distance to bank, and shelter density are suggested. For these and already measured variables, an examination of upper and lower tolerances for stage classes may be more useful from an ecological and conservation standpoint than examining in situ differences in means for the groups alone. Studies on larval Hellbender microhabitat during other seasons are also needed to determine if ontogenetic differences in microhabitat use occur throughout the year or are only limited to summer months.

Potential habitat differences among stage classes should be considered in future conservation and habitat restoration efforts, especially as accounting for multiple stage classes can assist in amphibian conservation efforts (Swanack et al, 2009). Immature individuals may be an important component for increasing some Hellbender population sizes as demonstrated by sensitivity analysis (Unger et al., 2013). Current Hellbender conservation efforts have focused heavily on head-starting and releasing individuals in order to boost adult populations. While these efforts are worthwhile and have proven successful (Bodinof et al., 2012a), consideration of immature Hellbender habitat at release and restoration sites is necessary to achieve the long-term goal of self-sustaining Hellbender populations. While related microhabitat needs may vary from site to site and should be studied in individual management areas, our study indicates that researchers and managers should consider heterogeneity in stream substrates, including fewer fine particles and more large gravel, in addition to a variety of boulders.

## ACKNOWLEDGEMENTS

Thank you to Dr. Marcy Souza, The Great Smoky Mountains Institute at Tremont, Dr. Perran Ross, Dr. Mary Christman, the Williams Lab at Purdue University, Andrea Drayer, Dr. Salvador Gezan, and all volunteers for assistance on this project. We would also like to acknowledge Paul Super, Keith Langdon, and the National Park Service. Financial support for this research was provided by the Great Smoky Mountains Conservation Association: Carlos C. Campbell Fellowship, The Reptile and Amphibian Conservation Corp (RACC), the Cryptobranchid Interest Group: Jennifer Elwood Conservation Grant. Research was conducted under permits from the National Park Service (GRSM-2009-SCI-0061, GRSM-20090056, GRSM-2008-SCI-0052, GRSM-00-131), and University of Florida ARC Protocol (#017-08WEC).

